# Microfluidic-based imaging of complete *C. elegans* larval development

**DOI:** 10.1101/2021.03.31.437890

**Authors:** Simon Berger, Silvan Spiri, Andrew deMello, Alex Hajnal

## Abstract

Several microfluidic-based methods for long-term *C. elegans* imaging have been introduced in recent years, allowing real-time observation of previously inaccessible processes. The existing methods either permit imaging across multiple larval stages without maintaining a stable worm orientation, or allow for very good immobilization but are only suitable for shorter experiments. Here, we present a novel microfluidic imaging method, which allows parallel live-imaging across multiple larval stages, while delivering excellent immobilization and maintaining worm orientation and identity over time. This is achieved by employing an array of microfluidic trap channels carefully tuned to maintain worms in a stable orientation, while allowing growth and molting to occur. Immobilization is supported by an active hydraulic valve, which presses worms onto the cover glass during image acquisition, with the animals remaining free for most of an experiment. Excellent quality images can be acquired of multiple worms in parallel, with little impact of the imaging method on worm viability or developmental timing. The capabilities of this methodology are demonstrated by observing the hypodermal seam cell divisions and, for the first time, the entire process of vulval development from induction to the end of morphogenesis. Moreover, we demonstrate RNAi on-chip, which allows for perturbation of dynamic developmental processes, such as basement membrane breaching during anchor cell invasion.

## Introduction

Traditionally, immobilization of *C. elegans* larvae and adults during image acquisition is accomplished using agar pads, a simple construct consisting of a glass slide and a thin slab of agar onto which individual worms are placed (Luke *et al.* 2014, Goodman *et al.* 1998). Immobilization is achieved by placing the cover glass on top of the agar pad and further improved using anesthetics such as levamisole or sodium azide (Sulston *et al.* 1977, Sulston *et al.* 1983). Whilst simple to fabricate, agar pad immobilization is invasive and results in a slowdown or complete arrest of development, precluding long-term observation (Fang-Yen *et al.* 2012, Wolke *et al.* 2007).

In recent years, a number of microfluidic approaches for *C. elegans* imaging have been reported (Chung *et al.* 2008, Mondal *et al.* 2016, Lee *et al.* 2014, Samara *et al.* 2010, Rohde *et al.* 2007, Hulme *et al.* 2010, Chronis *et al.* 2007). For example, *Gritti et al.* demonstrated a method in which worms are confined to large chambers (Gritti *et al.* 2016). Embryos placed in these chambers will develop and move freely, never leaving the region of interest set on the microscope. As animals are unrestrained, bodily motion is compensated by acquiring fluorescence images at high speed. Although broadly successful, the approach is limited to the assessment of bright fluorescent markers. More recently, *Keil et al.* presented a variation on this theme (Keil *et al.* 2017). Here, worms are confined to large microfluidic chambers and free to move for most of the experiment. Only during image acquisition are worms are immobilized; in this case using an inflatable hydraulic valve placed on top of the device that confines worms to the edge of the trap chamber upon actuation. As worms are immobilized in a secure fashion, a large variety of fluorescent markers and exposure times can be used. Such an approach is flexible, minimizes any negative effects on the animal and allows the study of development across all larval stages. Parallelization of experiments in either approach is achieved by fabricating multiple trap chambers on a single device, and sequentially imaging these throughout the experiment. However, as both of these approaches maintain worms in chambers significantly larger than the trapped worms, animals will move and rotate throughout the experiment, significantly hampering the tracking of many processes, and necessitating extensive post processing to compensate for animal motion. Lastly, Gokce *et al.* introduced a different approach for parallel long-term immobilization (Gokce *et al.* 2017). Here, a large number of worms are maintained in a large chamber for most of the experiment, allowing unhindered feeding and growth. During image acquisition, worms are pushed from the main chamber into an array of tapered channels, mechanically immobilized and imaged. Here parallelization is not achieved using multiple separate devices, but rather by using a number of parallel trap channels within a single device. Such an approach significantly simplifies device setup and operation, however repeatedly releasing worms from the trap channels into the large chamber does result in loss of animal identity as well as orientation. This in turn makes tracking developmental processes in the same worm impossible.

To address these limitations, we present a set of microfluidic devices that use “gentle” worm immobilization, therefore guaranteeing reliable development, whilst still allowing for high-resolution imaging and preservation of the orientation and identity of animals throughout an experiment. A first generation immobilization device, we introduced in 2018, allowed for the imaging of adult *C. elegans* under physiological conditions (Berger *et al.* 2018). Here, worms were trapped between two sets of on-chip hydraulic valves, confining the animal to the imaging area, while constantly supplying a bacterial food suspension and allowing egg laying through a set of pillars, placed next to a trapped animal’s vulva. Animals confined in this manner, on average remained viable for 100 hours, and could be imaged at high resolution using a variety of different imaging modalities. Furthermore, devices could be scaled to accommodate larvae at different stages, delivering the same advantages and performance as observed in adult *C. elegans.* However, this immobilization approach suffered from two drawbacks. First, parallelization of experiments was difficult due to the intricate channel system necessary to allow food supply and worm trapping, and second, *C. elegans* larvae could only be observed through a single larval stage at the end of which development arrested, as immobilized animals were unable to molt. We therefore developed a new generation of immobilization devices, which like the first generation devices preserve animal orientation and identity throughout an experiment and allow acquisition of high-resolution images, but allow imaging of multiple worms in parallel, as well as observation of worm development across multiple larval stages within a single device.

## Results

### Device Design and Function

Akin to the long-term immobilization approach proposed by Gokce *et al.* (Gokce *et al.* 2017), we performed long-term imaging using a parallel array of trap channels, rather than multiple individual trap chambers. In contrast to the approach by Gokce *et al.* (Gokce *et al.* 2017), we maintained the animals in the trap channels for the entire duration of the experiment, preserving worm identity and orientation. We developed three separate devices designed to cover all four larval stages. These devices are denominated as L1 for L1 larvae loaded right after hatching and imaged until the L2 stage; L1-L4 which can hold larvae from the mid to late L1 stage up to early/mid L4 stage; and lastly L2-A which can hold animals from approximately the mid L2 stage up to young adulthood. In all three devices, worms are trapped in up to 41 identical channels, which have a constant cross-section along the entire length, with only a slight taper at the head and a narrowing at the tail region of the channel (**Fig. 1a-a’,c**). Channel dimensions were chosen to be significantly longer and wider than the worms loaded at the beginning of an experiment, with a length/width of 400/15 μm for L1 devices, 575/20 μm for L1-L4 devices and 800/25 μm for L2-A devices respectively (**Fig. 1c**). In this way, worms have sufficient space to grow, as well as move to shed their skin during molting, but are prevented from turning or rotating. Channel height, unlike width, was chosen to be close to the thickness of the worms at the beginning of an experiment, i.e. 8 μm for early L1 larvae, 12 μm for mid/late L1 larvae and 15 μm for mid L2 larvae, in the L1, L1-L4 and L2-A devices, respectively. The reduced device height along with a carefully chosen device width keeps worms in a fixed orientation. Importantly, with a sufficiently small device height all worms are oriented and maintained in the desired lateral orientation. These stage-specific geometric constraints, necessary to prevent worms from rotating or turning head to tail, are the reason why three separate device types are needed to cover all larval stages. L1 larvae loaded into a channel large enough for young adult animals would simply not remain stable, and quickly leave the trap channel.

**Fig. 1.**
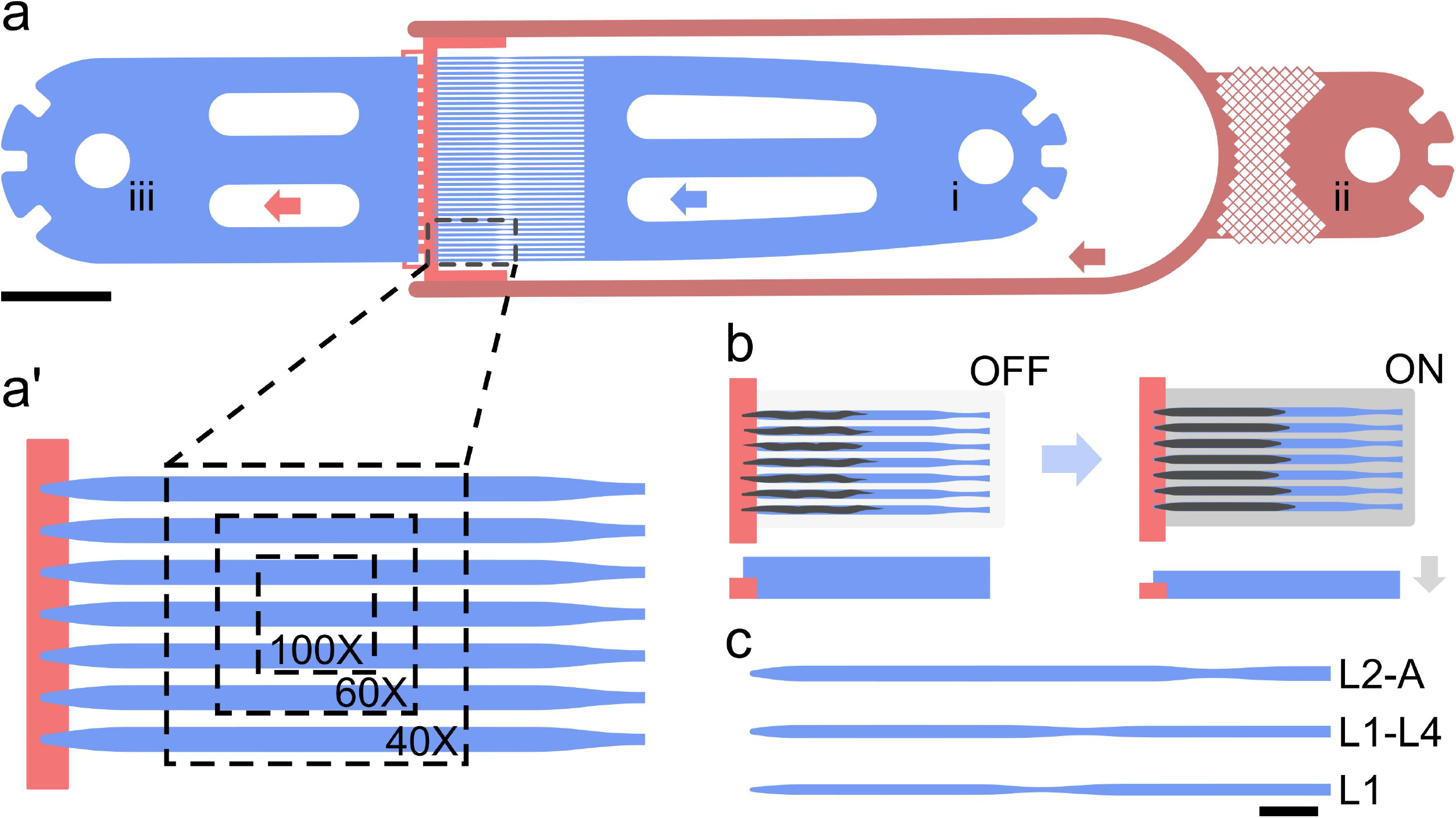
Microfluidic devices. (**a**) Device overview. Worms are loaded manually from (i) and pushed into a parallel channel array, housing up to 41 individual animals. A bacteria suspension is constantly supplied to all trap channels from (ii), with liquid leaving the device via the outlet (iii). Arrows indicate flow direction. (**a’**) Magnified view of the parallel imaging scheme. All trap channels (blue) connect to the same lower height food supply channel (red). Individual worms are held in the parallel channel array with device dimensions chosen to be significantly bigger than the worms initially loaded, allowing for growth over time. Bacterial food is constantly supplied via the low height channel (red) adjacent to the trap channel array. The low channel height allows for bacteria to flow through unimpeded and keeps worms from entering the food channel, effectively confining them to the imaging region of the device. Overlay shows the ROI achievable with different magnifications. (**b**) Functional principle. For most of the experiments, worms are held loosely in their respective trap channels with the active immobilization valve OFF. During image acquisition, the valve is turned ON with a pressure between 10 and 15 psi, resulting in the worms pressed onto the cover glass and immobilized. Once images are acquired, the valve is released. A side view of the trap channel is shown underneath. Channel height decreases in the ON state resulting in compressive force on the trapped animals (gray arrow indicates force direction). (**c**) Schematic of individual channels. Overview of the three device types and their relative scales. Scale bars are (**a**) 1000 μm and (**b-c**) 50 μm.

Food is supplied through an additional channel, which connects to the tip of each trap channel through a lower height section, i.e. a channel spanning the width of the device, with a significantly lower height than the trap region of the device (**Fig. 1a-a’**, red). The height of this connecting channel was chosen to be 4 μm for the L1 device and 5 μm for the L1-L4 and L2-A devices; large enough for bacteria to flow through unhindered and for the worms to reach the supplied food, but small enough to ensure that worms do not slip from the trap channel into the adjacent channel. Indeed, worms are trapped at the end of each channel not by a change in channel width, but by the sudden change in channel height at the transition from the trap channel to the food supply channel. All trapped worms passively remain at the tip of the trap channel within the designated imaging area (**Fig. 1a’**). L1 larvae were typically oriented in an oblique fashion, while L2 and older larvae remained on their lateral sides as on agar plates. It should be noted that whilst all 41 channels can be filled simultaneously with animals, not all worms will be facing the food supply channel. Only worms facing the food supply channel will be able to feed and develop normally, while worms facing away from the food supply will develop at a slower rate, especially toward the end of an experiment. Care must therefore be taken to load worms in a “head first” orientation, through careful manipulation during device loading. In the experiments presented herein, a total of 187 worms were imaged. Of these 159 (85%) were oriented correctly with the head facing the food source and only 28 (15%) were oriented incorrectly.

As worms are maintained in channels that are too large for passive immobilization, additional active immobilization measures were implemented using a large inflatable hydraulic valve fabricated on top of the array of parallel trap channels. Inflating this hydraulic valve results in a downward force that presses trapped animals onto the cover glass and minimizing animal motion (**Fig. 1b**). The hydraulic valve is only inflated during image acquisition, such that worms remain nearly free for most of the experiment. It should, however, be noted that even when the hydraulic valve is actuated animals are not completely immobile. This may be remedied by applying higher pressure, Though, we found that application of excessive pressure negatively affects developmental speed. Nonetheless, worms trapped in our devices and compressed with a pressure of 10-15psi remained stable enough for acquisition of high-resolution images, suitable for long-term tracking and 3D reconstruction of developing features of interest (**Movie S1**). Residual motion is essentially linear along the length of the trap channel (**Movie S2**) with features of interest shifting as worms grow in size. This linear motion can be compensated for through automated or manual image registration. This greatly simplifies the tracking of processes, especially when compared to other microfluidic methods, which deal with large shifts in animal position and require computational straightening of features of interest. Image quality is unaffected by the microfluidic device itself, since these are fabricated on a cover glass and made from an optically transparent elastomer. Indeed, devices are inherently compatible with high magnification, high NA immersion objectives, with no noticeable deterioration in image quality

Parallelization is achieved by spacing the trap channels such that multiple animals can be imaged at the same time. Channel spacing was chosen such that in the field of view of a sCMOS camera with a 18.7mm diagonal chip area, at 100X magnification three, at 60X five and at 40X seven animals can be simultaneously imaged in a single region of interest (**Fig. 1a’**). Throughput can be further increased using a motorized XY-stage to acquire multiple regions of interest within the same microfluidic device, generating a throughput equal to or higher than other published systems.

### Development On-Chip

On-chip developmental timing was established by following the seam cell divisions, which occur at the beginning of each larval stage, using the three separate devices for L1, L1-L4 and L2-adulthood (**Fig. 2**). The hypodermis was visualized using a DLG-1::GFP marker (*dlg-1(mc103[dlg-1::gfp]))* (Vuong-Brender *et al.* 2017) in experiments with the L1 device and a HMR-1::GFP adherens junction marker (*cp21*) (Marston *et al.* 2016) in experiments with all other devices. Both markers outline all seam cells as well as the P-cells and at later stages also the vulval precursor cells (VPCs). The DLG-1::GFP marker was used in experiments with L1 larvae since HMR-1::GFP shows a very low signal to noise ratio at this stage, making it difficult to identify individual cells with this marker.

**Fig. 2.**
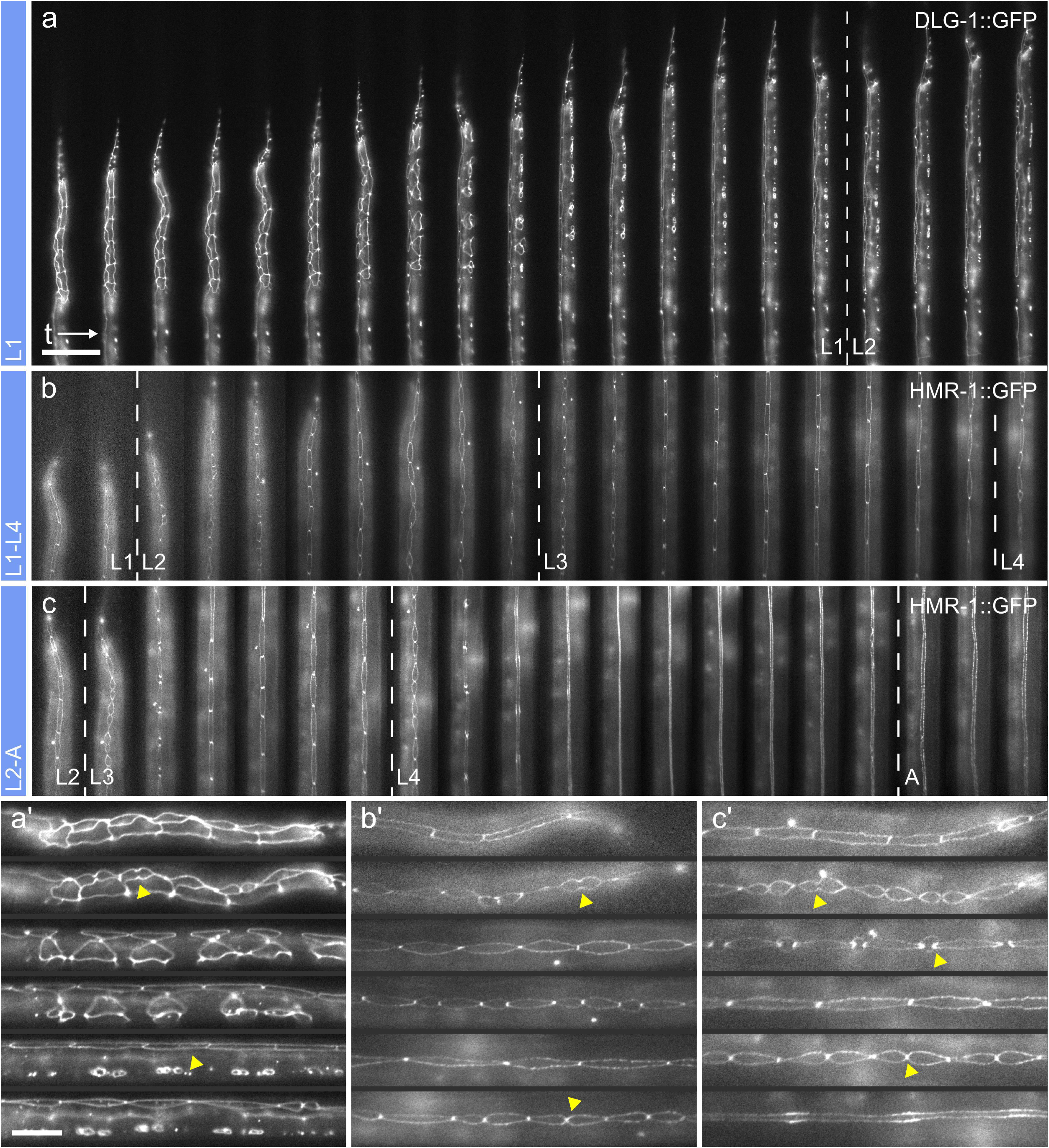
Developmental Timing. Reliability of development on-chip was assessed by imaging development of the hypodermis across all larval stages. (**a**) Single L1 stage larva imaged from starvation-induced arrest after hatching until the L2 stage using the L1 device. Both seam cell and P-cell divisions are shown. The dashed line indicates the transition between L1 and L2, determined by the first seam cell division in L2. (**b-c**) Cropped views of a (**b**) single larva imaged from late L1 across L2 and L3 until the L4 stage using the L1-L4 device, and from the (**c**) a late L2 stage, imaged across the L3 and L4 stages until adulthood using the L1-L4 device. As in L1 larvae, development was scored using the hypodermis, with the occurrence of the first seam cell division indicating the end of the previous and beginning of the next larval stage or in L4, the occurrence of vulval eversion indicating the beginning of adulthood. Dashed lines indicate transitions between stages. (**a’-c’**) Magnified view of selected time points for images shown in (**a-c**) respectively. Highlighted are some of the various seam cell division and fusion events observed along the worm development. Time progression from left to right and time frames are shown at 1-hour intervals. Raw images acquired at 40X magnification are shown. Scale bar in (**a**) 50 μm and in (**a’**) 20 μm, same scale for (**b-c**) and (**b’-c’**) respectively.

Developmental timing for the L1 devices was assessed from the start of an experiment (time zero) up to the L2 stage. Several milestones were assessed in timing and reproducibility studies. These included the time at which all seam cells had finished their respective divisions and reconnected to their neighbors, the end of all ventral P-cell migrations and divisions in the L1 stage, as well as the time at which the first seam cell divided in the L2 stage (**Fig. 2a**). We found that on average, all seam cells had divided and reconnected to their neighbors after 12.0±1.7 hours in L1 (*n* = 22), and P-cells had concluded their ventral migration and divided along the A-P axis after 14.9±1.4 hours (*n* = 22). The first seam cell division in L2 was typically observed after 21.16±3.5 hours (*n* = 20) and completion of all L2 divisions after another 12.9±4.0 hours (*n* = 12). These data indicated that the time in which these processes occur is highly reproducible. Developmental timing, especially early on during development, i.e. seam cell divisions and fusion to the hypodermis occur at timing consistent with literature values (Soulston *et al.* 1983). However, the start of seam cell division in the L2 stage appeared to be delayed and more variable. This is most likely caused by the small size of the trap channel needed to properly orient early L1 larvae at the start of the recording, with the narrow confinement most probably impeding the L1/L2 molting. Nonetheless, using the L1 device all imaged animals developed through the L1 stage. Whilst not recommended due to possible molting delays, development in the early L2 stage can still be assessed using this device.

For the L1-L4 and L2-A devices, we likewise quantified developmental timing using the time at which the first seam cell initiates a new round of division indicating the end of the previous larval stage and the start of a new one. The length of L2 and L3 were 10.2±1.0 and 10.9±1.4 hours, respectively (*n* = 27) using the L1-L4 device (**Fig. 2b**), and for L3 and L4 the time was 10.4±2.3 and 14.6±2.2 hours respectively (*n* = 19), using the L2-A device (**Fig. 2c**). The end of L4 stage was not defined as the moment when seam cells fuse (which happened on average after 5.7±1.3 hours, *n* = 19), but as the time when vulval eversion is completed, indicating the end of vulval development. All animals imaged in the L1-L4 and L2-A devices reached the L4 stage and adulthood, respectively. For the L2-A device it should be noted that development does not immediately arrest in early adulthood, with adult worms remaining viable for several more hours, with multiple ovulation events being observed in trapped animals.

Comparison to worms grown on NGM plates under standard conditions yielded median developmental times for L1 15 hours (*n* = 21), L2 10 hours (*n* = 46) for the DLG-1::GFP strain, for the HMR-1::GFP strain we found the following values L2 10 hours (*n* = 15), L3 12 hours (*n* = 21) and L4 12.5 hours (*n* = 61) (**Fig. S1**). These data indicate that development on NGM plates occurs slightly faster than on-chip, especially in L1 larvae and during the L4 stage. This is most likely caused by the low channel height, which over time compresses the growing worms and results in a delay in developmental speed.

### Post-embryonic development of the hypodermis

Using the smallest (L1) of our three devices, we studied the development of the hypodermis, specifically assessing seam cell and P-cell division and migration. Our aim was to gain insights into the variability of developmental timing, which cannot be accessed by imaging single animals on agar pads. Seam cells divide in a fixed pattern during all four larval stages. Their divisions are asymmetric, with the posterior of the two daughter cells normally remaining as a seam stem cell and the anterior fusing with the surrounding hypodermis (**Fig. 3a)** (Hedgecock *et al.* 1985). The P-cells are initially connected to neighboring seam cells, forming pairs along the A-P axis (Hedgecock *et al.* 1987, Sulson *et al.* 1977). Over the course of the L1 stage, the P-cells detach from the seam cells, migrate towards the ventral midline and divide, with the posterior daughter cells of P1, P2 and P9-P11 (the Pn.p cells) fusing to hyp7 and the anterior Pn.a cells differentiating into ventral cord motor neurons. The central Pn.p cells P3-8.p remain unfused and become the VPCs (except for P3.p, which fuses later during L2 in around 50% of the cases) (Sulston *et al.* 1983).

**Fig. 3.**
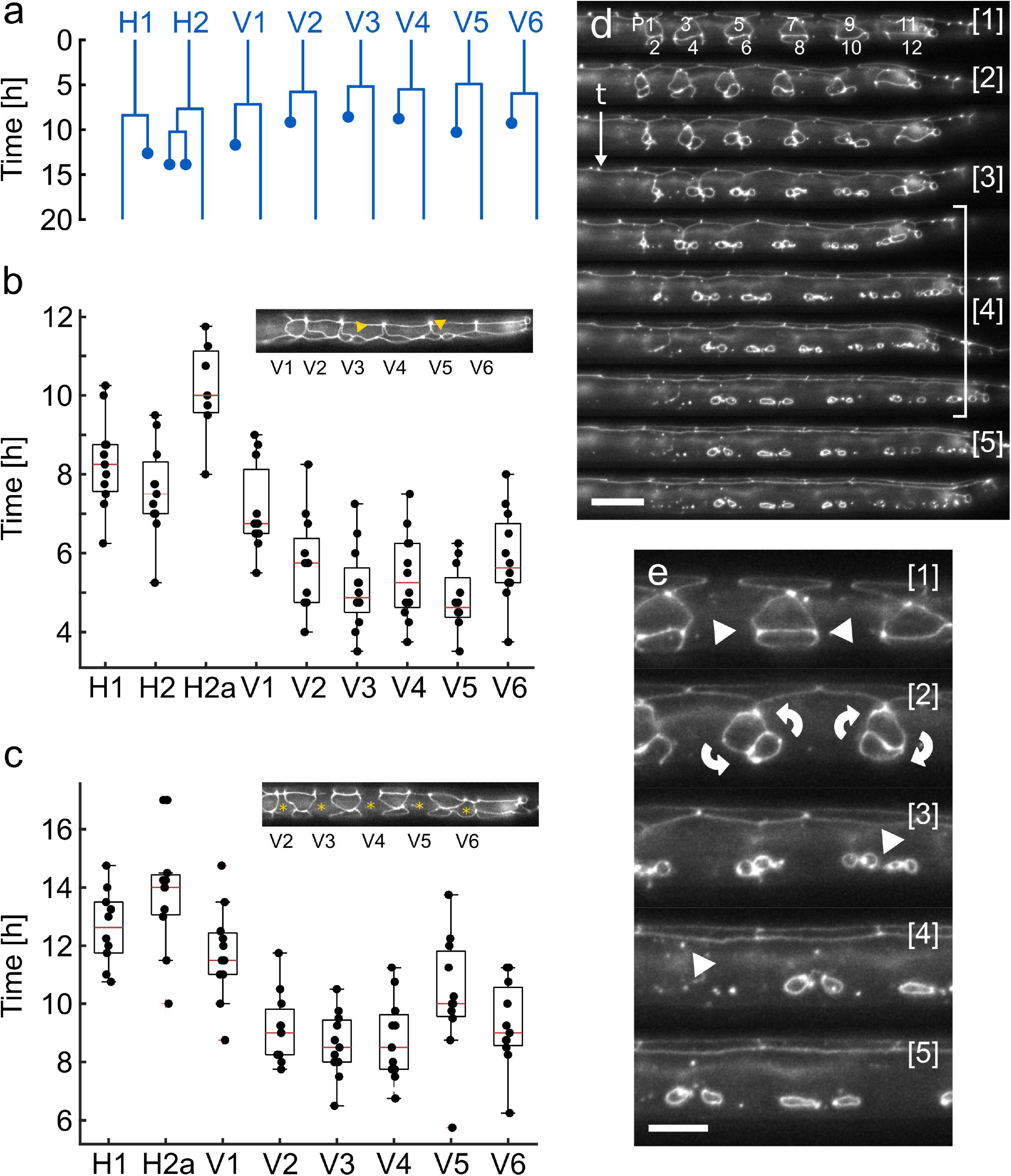
Development of the Hypodermis. (**a**) Average seam cell lineage constructed from all L1 larvae analyzed. The seam cells except for H0, which does not divide in any larval stage, and the posterior T cell and its descendants are shown. The average time by which each cell divides is indicated, with V5 dividing first and H1 dividing last. The times at which one or several of the daughter cells fuse with the surrounding hyp7 are represented by the dots. (**b**) Quantification of the seam cell divisions across the L1 stage (*n* = 12). Insert: representative image showing V3 and V5 post division with all other seam cells still undivided. (**c**) Quantification of the time at which the anterior daughter seam cells fuse with hyp7 (*n* = 12). Box plots in (**b**) and (**c**) show the median (red lines) with upper and lower quartiles, bars indicate the most extreme points of the distribution. Data in (**b**) and (**c**) were used to derive the average seam cell lineage shown in (**a**). Insets show representative images in which V4a has completely fused with hyp7, V3 and V5 are close to complete fusion and all other seam cells are still unfused. (**d**) Representative images showing P-cells migrating from their initial location toward the ventral midline, as well as the divisions along the A-P axis and cell fusion manifesting by the dissolution of the HMR-1::GFP labelled cell junctions. [1] shows the six pairs of P-cells (*, from left to right P1/2 through 11/12) initially attached to V2a-V6a. [2] P-cells begin migrating and rotating toward the ventral midline. [3] P-cells in mid migration, with some already aligned with the ventral midline. [4] P-cells divide and some descendants fuse with the surrounding hypodermis. [5] Six P-cells (P3.p-P8.p) that become the VPCs remain unfused and will divide in L3 to form the vulva. Time progression from top to bottom, images are shown at 45-minute intervals. (**e**) Magnified views of time points [1-5] shown in (**d**). Raw images acquired at 40X magnification are shown. Scale bar (**d**) 15 μm and (**e**) 7.5 μm.

Using our L1 device, we were able to image worms loaded immediately after hatching and until the early L2 stage. Images were acquired at 15-minute intervals, which are short enough to observe all individual cell divisions and long enough for imaging not to affect normal development. We were able to identify and follow all seam cells (**Fig. 3a-c, Movie S2**), except for those originating from the tail cell, which were typically obstructed in the oblique orientation of L1 larvae. All seam cells imaged on-chip followed the same pattern of asymmetric cell divisions, indicating that the microfluidic device does not affect the process (*n* = 12). As previously described, the V5 cell is usually the first seam cell to divide, followed by V3 with H1 and H2 dividing last (**Fig. 3b**). A similar pattern was observed for the times at which daughter seam cells fuse with hyp7, except that V3 and V4 were often the first cells to fuse even though they are not necessarily the first ones to divide (**Fig. 3c**). Similarly, we could identify all 12 P-cells and follow their migration toward the ventral midline and divisions into the Pn.a and Pn.p cells (**Fig. 3d**). Before migrating ventrally, the pairs of P-cells rotated by 90° to align along the A-P axis. On average, P-cells began division after 11.7±1.6 hours (*n* = 12) and the fusions of their daughter cells was completed after 15.9±2.7 hours (*n* = 12). Importantly, images acquired on chip were of excellent quality, allowing identification and tracking of all GFP-labelled cells of interest in 3D reconstructions (**Movie S3**).

### Observing VPC induction

The second device type, L1-L4, was tested by observing vulval induction. The development of the hermaphrodite vulva is one of the best-studied models for organogenesis, as many mutants have been isolated that affect various aspects of the process (Sternberg *et al.* 1989, Sternberg *et al.* 1986, Sternberg *et al.* 2005, Greenwald *et al.* 1983). During the L2 stage, three of the six VPCs (P5.p, P6.p & P7.p) are induced to adopt one of two vulval cell fates (1° or 2° fate). VPC fate specification occurs through a combination of an inductive epidermal growth factor (EGF) signal, secreted by the gonadal anchor cell (AC), together with lateral Delta/Notch signaling among the VPCs. In wild type animals, the AC is situated closest to P6.p. Thus, P6.p receives the highest amount of inductive EGF signal, which specifies the 1° fate by activating the EGFR/RAS/MAPK signaling pathway. The adjacent VPCs P5.p and P7.p receive less inductive signal and acquire the 2° vulval fate due to the activation of the lateral Notch signaling pathway. The three remaining VPCs (P3.p, P4.p and P8.p) receive only little inductive and lateral signal and adopt the non-vulval 3° fate by dividing once and then fusing with hyp7. Following its induction, P6.p divides thrice to generate 8 descendants, while its neighbors P5.p and P7.p generate 7 descendants in an asymmetric lineage. After the three rounds of vulval cell divisions have been completed by the end of the L3/beginning of the L4 stage, a total of 22 vulval cells have been generated.

We observed the specification of the 1° cell fate using the *arIs92[egl-17::cfp]* transcriptional reporter (**Fig. 4a**, **Movie S4)** (Yoo *et al.* 2004). In animals recorded from the late L1 stage onwards, weak EGL-17::CFP expression first became apparent during the early L2 stage (**Fig. 4a-b**). At this stage, EGL-17::CFP-fluorescence could not only be observed in P6.p, but also in the neighboring VPCs P5.p and P7.p (**Fig. 4a-b**). Only towards the end of the L2 stage, when P6.p has irreversibly been determined as the 1° VPC, the other VPCs ceased expression of the 1° fate marker (**Fig. 4a-b**). Once the 1° fate had been established, EGL-17::CFP expression further increased in P6.p, and during the subsequent VPC divisions EGL-17::CFP remained highly expressed in the P6.p descendants (**Fig.4 a,c**). Along with EGL-17::CFP fluorescence, we quantified gonad length as an accurate measure of the developmental stage (**Fig.4 d**) (Mereu *et al* 2020). Especially during the L2 and early L3 stages, gonad length increased linearly and relatively uniformly in the imaged worm population (*n* = 19), indicating reliable on-chip development. The variation in gonad length increased toward the end of the L3 stage, as part of the gonads became obscured by the intestine. EGL-17::CFP fluorescence intensity in P6.p and its descendants was quantified for each worm over time, revealing a relatively slow intensity increase throughout the L2 stage, followed by a more rapid increase during the L3 stage. Interestingly, the EGL-17::CFP signals varied greatly from animal to animal, even after the 1° fate had been established in late L2 (**Fig. 4e**), reflecting the dynamic and initially variable nature of vulval induction within an isogenic population of animals.

**Fig. 4.**
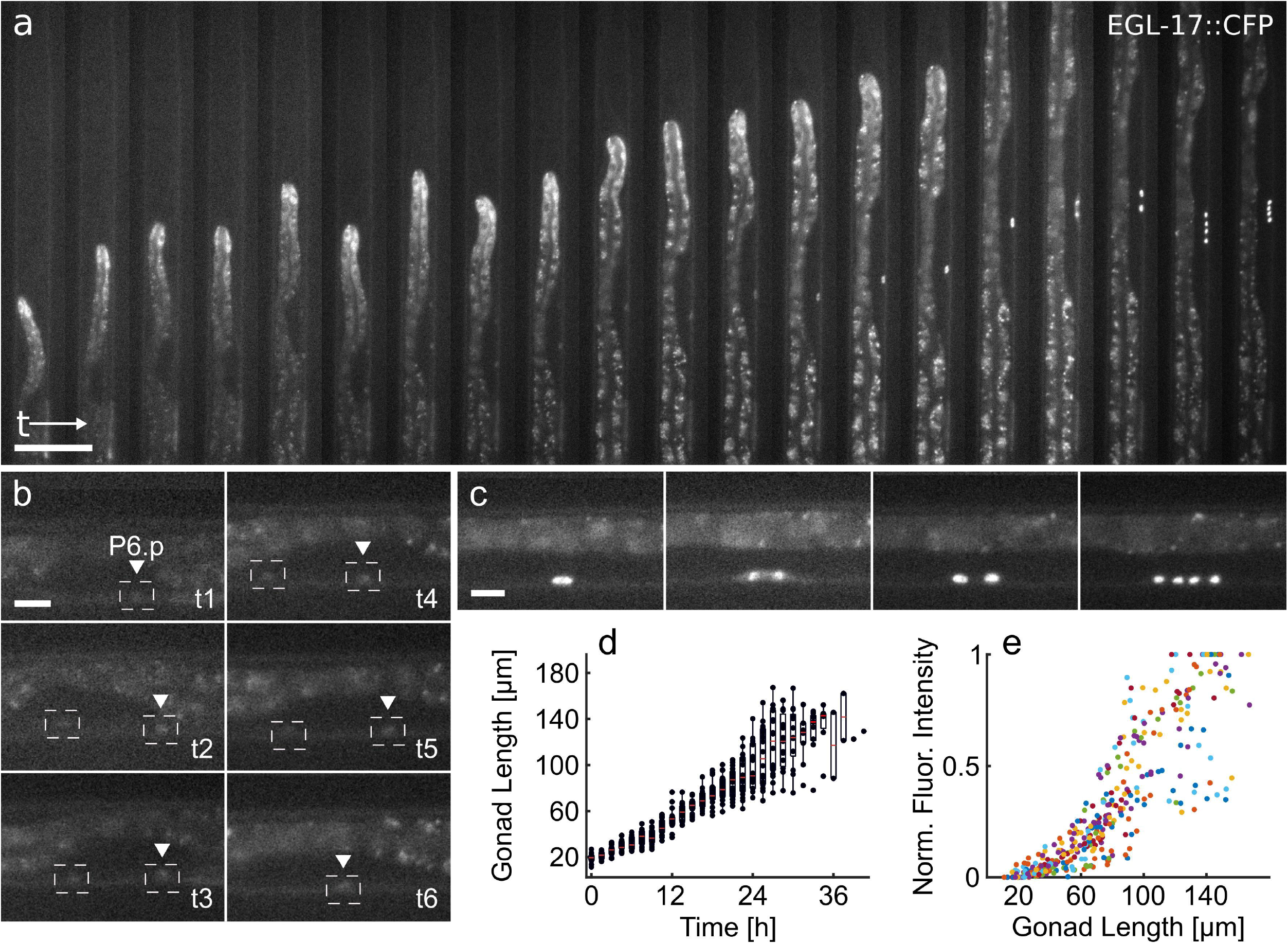
Vulval fate specification. (**a**) Overview of vulva induction from late L1 to the 4-cell stage in mid L3. Stable expression of the 1° VPC fate reporter EGL-17::CFP in P6.p is established over the course of the L2 stage, followed by three rounds of cell division. The first two rounds of VPC division were recorded. Time progresses from left to right at 1.5-hour intervals. (**b**) Magnified views of the worm in (**a**). EGL-17::CFP expression first becomes apparent in P6.p during the early L2 stage (t1), but initially expression is also detectable in P7.p (t2-t5), until the 1° fate is firmly established in P6.p during L2 (t6). (**c**) Magnified views of the worm in (**a**) during L3 highlighting the 1° fated VPC (P6.p) before its first division, during the first division, as well as after the first and second divisions. (**d**) Gonad length measured over time (*n* = 19). Especially during the L2 and early L3 stages gonad length increases linearly with time, showing little variation between animals. Measurement variation increases toward the end of L3 stage as the gonad becomes partially obstructed by the intestine. The linear increase in gonad length especially early on during the experiment indicates reliable development on-chip. (**e**) Normalized maximum EGL-17::CFP fluorescence intensity relative to gonad length as timing reference (*n* = 19). The observed fluorescence intensities vary substantially between individual worms, which are indicated by color-coded dots. The box plots in (**d**) show the median values (red lines) with upper and lower quartiles and bars indicating the extremes of the distribution. Raw images acquired at 40X magnification. Scale bars are (**a**) 50 μm and (**b-c**) 10 μm.

In summary, these experiments show the suitability of our L1-L4 device in imaging VPC induction and the possibility of quantifying gene expression levels based on fluorescent reporters at high temporal resolution.

### Live-imaging vulval morphogenesis

The capabilities of the third set of devices (L2-A), were assessed by imaging the entire process of vulval development until the completion of vulval morphogenesis (Schmid *et al.* 2015, Schindler *et al.* 2013). As with the seam cell divisions, the formation of the vulva occurs in a stereotypical fashion with the formation of different structures that can be used to track the reliability and test for normal developmental progression. After the induction of the three VPCs at the late L2/early L3 stage, three rounds of cell divisions yield a total of 22 vulval cells by the end of the L3 stage. During the L4 stage, epithelial morphogenesis gives rise to the complete vulva. The 22 vulval cells invaginate to form a ventral lumen and extend circumferential processes that fuse with their contralateral partner cells, thereby forming a stack of seven toroids that build the tubular organ. Finally, by the end of the L4 stage, the vulval toroids are connected to the uterus and eversion of the vulva closes the lumen.

We imaged vulval development from the early L3 stage onwards, visualizing all cell divisions and the entire process of morphogenesis using the HMR-1::GFP adherens junction marker described above. Additionally, the AC was labelled with the *Pcdh-3>mCherry::moeABD (qyIs50)* reporter, as the AC plays key roles during vulval induction and morphogenesis. Using these two markers, we were able to visualize the divisions, migrations, invasion, fusions and shape changes of the 22 vulval cells during the L3 and L4 stages with excellent spatial and temporal resolution, allowing us to obtain 3D reconstructions of the process (**Fig. 5a-a’’, Movie S5**). As with the other processes studied, we ensured that imaging in the microfluidic device does not affect development in morphology or speed. To this end, we assessed the time needed to progress from invagination (L4.0, *t1)* to the mid L4 stage (L4.5, *t4*) according to the sub-stages defined by Mok et al. (Mok *et al.* 2015) (**Fig. 5b-c**). The average progression time between *t1-t2* was 3.8±1.2 hours, between *t2-t3* 3.5±1.2 hours and between *t3-t4* 2.8±1.6 hours (*n* = 7), indicating an approximately linear developmental progression (**Fig. 5c**). Like in the developmental timing assay these times, especially *t1-t2,* appear slightly delayed when compared to literature values (Mok *et al.* 2015), which may stem from the narrow dimensions of the device delaying molting. Nonetheless, developmental timing within the animals we assessed is highly reproducible and all animals completed development into adulthood. We then proceeded to measure multiple features of the toroids at selected time points (M1-M5 in **Fig. 5b**), with the variation in measurement again serving as an indication of developmental consistency. The measurements of the selected features exhibited little variation between animals (**Fig. 5d,** M1 7.64±0.54 μm; M2 4.39±0.56 μm; M3 6.02±0.45 μm; M4 13.80±0.56 μm; M5 54.71±4.36 μm, *n* = 7). Together, these measurements indicate that our microfluidic devices do not significantly affect developmental patterns or reliability. Furthermore, as shown in **Fig. 5a-b** our microfluidic imaging method combined with standard image deconvolution methods delivers high image quality, enabling us to identify and track individual cells in 3D reconstructions throughout the entire developmental process (**Movie S5**). In combination with the excellent temporal resolution, this method has allowed us to visualize for the first time vulval morphogenesis in its entirety, uninterrupted across the L3/L4 molt.

**Fig. 5.**
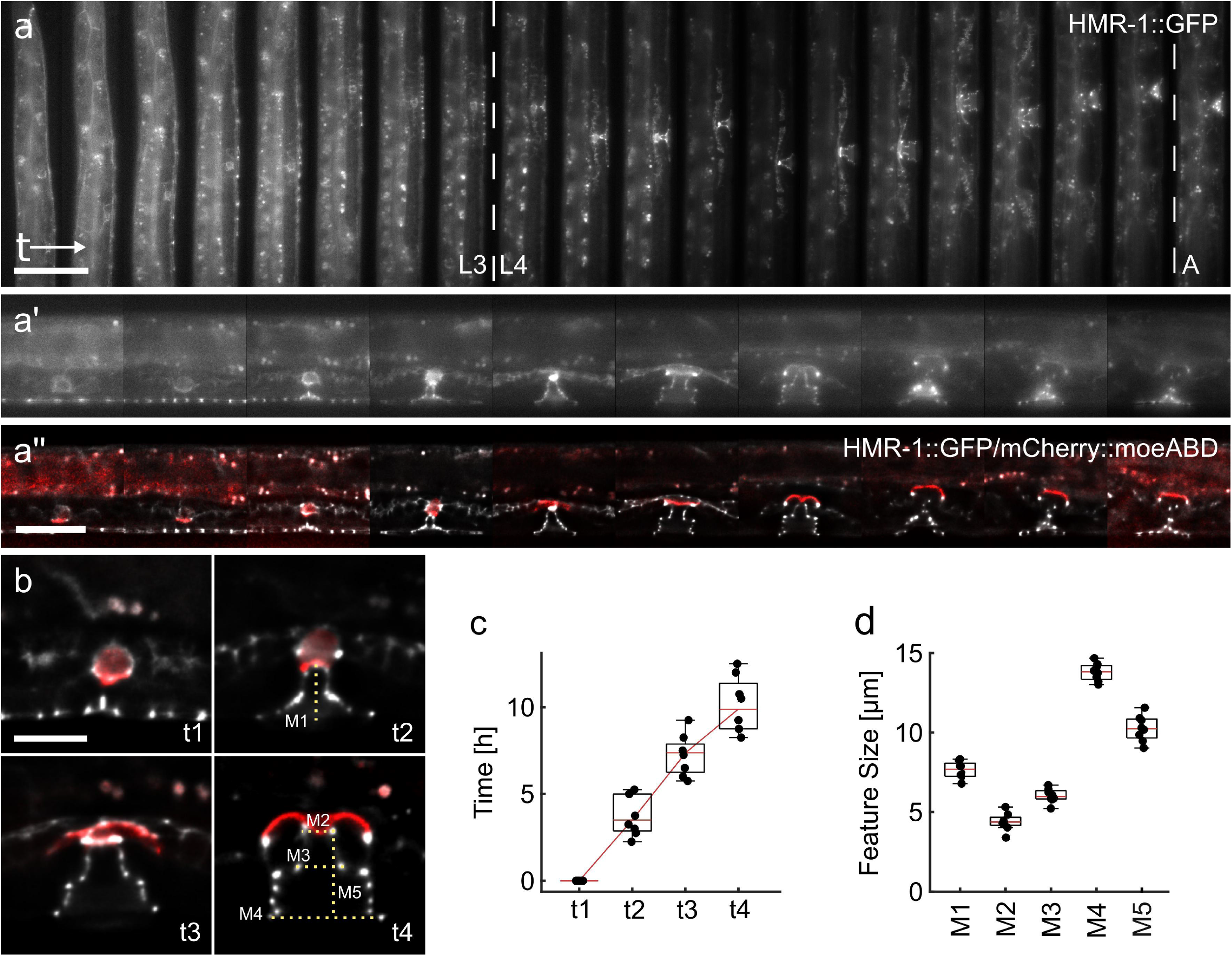
Vulval morphogenesis. (**a**) Overview of complete vulval development. The HMR-1::GFP marker was used to observe the cell junctions of the VPCs and the toroids forming the vulva. A single animal, with time progressing from left to right at 1-hour intervals shows the three rounds of VPC divisions during L3, followed by vulval invagination and toroid formation in L4 and finally eversion of the vulva during L4/adult transition. (**a’**) Magnified views of selected time points from (**a**), illustrating the excellent raw image quality attainable. (**a’’**) Magnified and deconvolved z-sections of the same frames shown in (**a’**) illustrating the attainable image quality. The HMR-1::GFP signal is overlaid with the mCherry::moeABD marker in red outlining the AC (**Movie S5)**. (**b**) Vulval development at the L4.0 (t1), L4.1 (t2), L4.3 (t3) and L4.5 (t4) stages defined by (Mok *et al.* 2015). The average time needed for animals to transition from one time point to the next, as well as morphological features indicated by the dotted lines and M1-5 were measured. (**c**) Time needed for animals to develop from L4.0 to L4.5, indicating a linear transition through different developmental checkpoints and little variation between animals. (**d**) Sizes of the features M1-M5 measured for all animals, indicating close clustering of measured values. The box plots in (**c**) and (**d**) show the median (red lines) with upper and lower quartiles and bars indicating the extremes of the distribution. Raw images acquired at 60X magnification, except for (**a’’**) and (**b**), which show deconvolved z-sections. Scale bars are (**a**) 50 μm, (**a’**) 25 μm and (**b**) 10 μm, same scale for (**a’**) and (**a’’**).

### On-Chip RNAi Perturbation of AC Invasion

As a final case study, we assessed whether our microfluidic devices could be used to study perturbation of developmental processes *in vivo.* For this purpose, we inhibited basement membrane (BM) breaching during AC invasion (Sherwood *et al.* 2003) by RNAi-mediated inhibition of the *egl-43* gene, which encodes an essential transcription factor controlling AC invasion and proliferation (Hwang *et al.* 2007, Rimann *et al.* 2007, Deng *et al.* 2020, Sherwood *et al.* 2005). In wild-type animals, AC invasion invariably occurs during the late L3 stage, resulting in the dissolution of the BMs separating the uterus from the vulval epithelium and the establishment of a direct connection between the AC and 1° vulval cells (Sherwood *et al.* 2003). RNAi against *egl-43* by feeding worms dsRNA producing *E.coli* bacteria on plate causes an up to 96% penetrant BM breaching defect (Deng *et al.* 2020). Accordingly, we fed worms on-chip with *E.coli* carrying either an empty vector control plasmid or expressing *egl-43* dsRNA. In worms grown on empty vector containing bacteria, all but one of the animals imaged displayed normal BM breaching by the end of the L3 stage (*n* = 25) (**Fig. 6a-a’, Movie S6**). By contrast, in worms fed on chip with *egl-43* RNAi bacteria, none of the L3 animals imaged showed any signs of BM breaching, with the BM remaining intact well into the L4 stage (*n* = 16) (**Fig. 6b-b’, Movie S7**). Furthermore, a mispositioning of the AC as well as AC proliferation was often observed (**Movie S7**), as reported previously (Deng *et al.* 2020). These experiments demonstrated that our microfluidic devices are well suited for *in vivo* perturbation experiments.

**Fig. 6.**
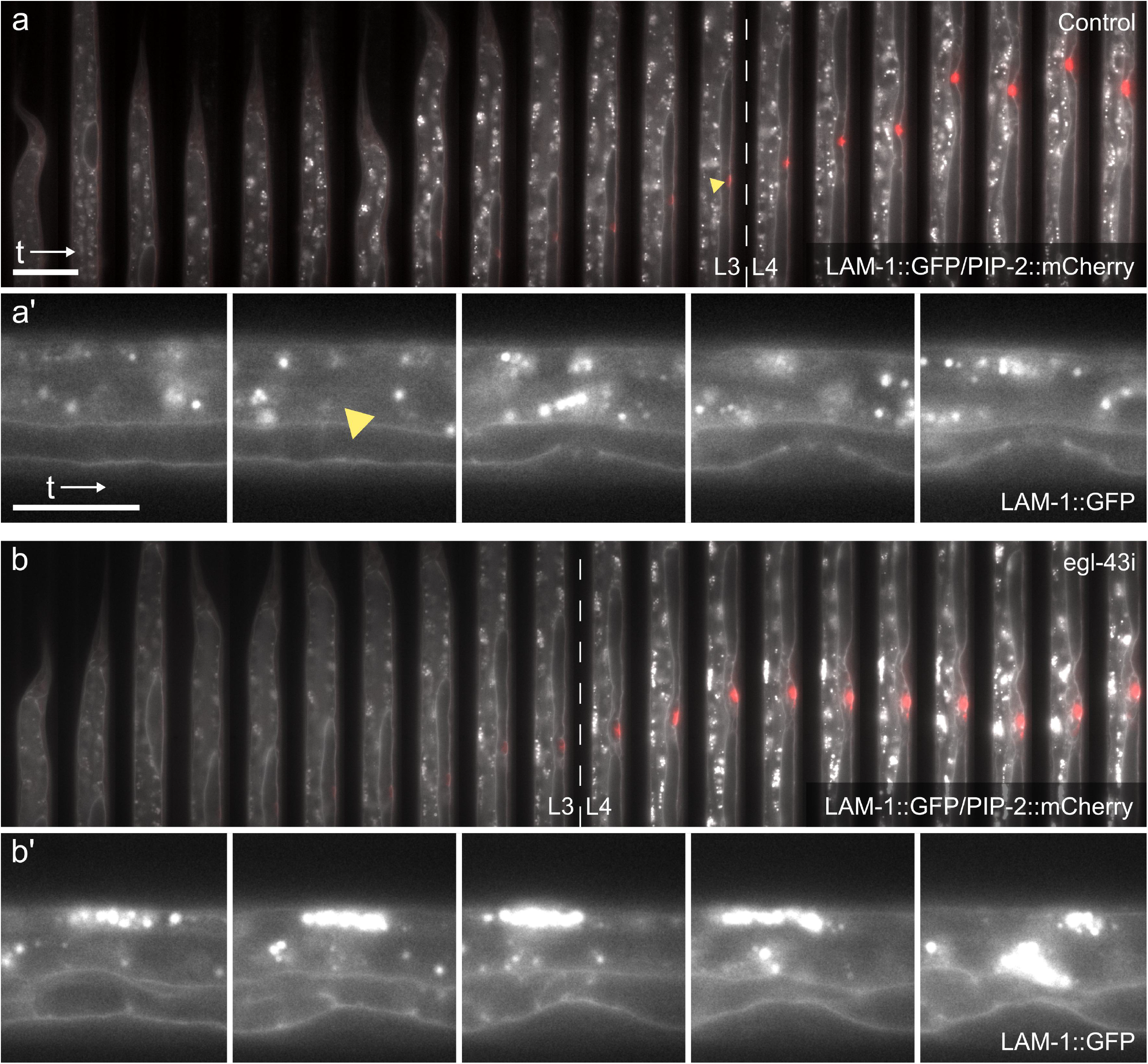
On-chip perturbation of AC invasion. (**a**) Overview of AC invasion in an empty vector treated control. A single worm imaged from L2 until late L4 is shown, progressing from left to right at 1-hour intervals. The LAM-1::GFP (*qyIs10[lam-1::gfp; unc-119 (+)] IV*) marker labelling the BMs (gray) and the *Pcdh-3>mCherry::moeABD* reporter(*qyIs50*) outlining the AC (red) were used. (**a’**) Magnified views of time points selected from (**a**). During late L3, the BMs are breached (yellow arrowheads), shortly before vulval invagination begins. (**b**) AC invasion in an *egl-43* RNAi-treated animal. Contrary to the control shown in (**a**), the BMs are not breached and remain intact after invagination well into the L4 stage. The AC is misaligned with the 1° VPCs and multiple ACs are formed during L4. (**b’**) Magnified views of time points selected from (**b**). Raw images acquired at 60X magnification. Scale bars (**a**) 50 μm and (**a’**) 25 μm, (**b**) and (**b’**) are shown at the same respective scales.

## Discussion

Herein, we present a novel microfluidic long-term imaging platform that allows for high-resolution imaging and tracking of a variety of developmental processes in *C. elegans* during all four larval stages up to adulthood. Moreover, our devices permit the imaging of multiple worms in parallel and across different larval stages, as animals on-chip can undergo molting and transition from one to the next larval stage. Contrary to previously published microfluidic-based longterm imaging strategies, this novel approach allows the preservation of both worm identity and worm orientation throughout the experimental timeframe, facilitating tracking of complex developmental processes in time. Trapping of up to 41 individual worms is accomplished using an array of identical trap channels, connected to an on-chip food supply and covered by a single large hydraulic valve. Immobilization of all trapped worms is achieved through the periodic inflation of a hydraulic valve, resulting in the trapped worms being pressed onto the cover glass only during image acquisition. High throughput imaging can be achieved through the narrowly spaced trap channel array, in combination with the large sensor of modern sCMOS cameras, simultaneously imaging between 3 and 14 animals, depending on the chosen magnification.

Device function, specifically whether it allows development under physiological conditions, was assessed in all four larval stages. Seam cell divisions during different larval stages were imaged using all device types and compared to the approximate developmental timing observed in freely crawling worms grown on NGM plates. We found very consistent development on-chip as evident from the low standard deviation of division times in each stage, as well as the generally good agreement between developmental timing on-chip and on NGM plates. Any observed differences most likely result from prolonged confinement through the low height of the device channels, as deviation from normal timing typically becomes pronounced during later stages when animals outgrow a specific device. These findings are further supported by the reliable on-chip development observed in all other processes studied, such as P-cell migration, VPC induction, AC invasion and vulval morphogenesis. Using these devices, we were therefore able to gather data on developmental timing previously not available, for example by following both individual seam cells and P-cells throughout L1, vulval precursor cell induction and divisions or toroid formation during L4. In particular, we were for the first time able to observe the entirety of vulval morphogenesis in individual animals across the L3/L4 molt and obtain 3D reconstructions of the developing organ. This approach will be yielding new insights into the mechanism controlling vulval morphogenesis. Finally, we have demonstrated that our devices are not only suited to study wild-type *C. elegans* development, but can also be employed to perform gene perturbations by RNAi. On-chip perturbation may also be applied using other systems, such as using small molecule inhibitors or the auxin-inducible protein degradation system to inhibit proteins of interest (Zhang *et al.* 2015). These approaches will become invaluable when investigating the dynamic regulation of developmental processes.

Taken together, our results demonstrate the utility of a novel long-term imaging approach in studying a variety of developmental processes during *C. elegans* larva development with excellent image quality and without significant negative impact on developmental timing.

## Materials and Methods

### Worm Maintenance and Preparation for Microfluidic Imaging

Worms were maintained at 20°C on Nematode Growth Medium (NGM) plates seeded with *E. coli* OP50 as described (Brenner 1974) Prior to an experiment, worms were synchronized by bleaching of gravid adult hermaphrodites (Stiernagel 2006). The collected embryos were pelleted by centrifugation (1300rcf for 1 minute) and washed with fresh S-Basal buffer twice. Embryos were left to hatch in S-Basal overnight, arresting larvae in the L1 stage. After overnight incubation, arrested larvae were passed through a 10 μm filter (pluriStrainer Mini 10 μm, PluriSelect, USA), removing debris as well as unhatched embryos, washed twice with fresh buffer and grown on NGM plates. Once the synchronized population had reached the desired stage, worms were washed off the NGM plates using S-Basal buffer, washed twice with fresh S-Basal to remove any debris and loaded into the device. Larvae used in experiments with L1 animals were not grown on NGM plates but directly loaded into the device right after washing. The following alleles and transgenes were used in this study:

LGI: *hmr-1(cp21[hmr-1::gfp + LoxP]*) (Marston *et al.* 2016).
LGII: *arIs92[egl-17::cfp]* (Yoo *et al.* 2004).
LGII: *qyIs23[cdh-3>mCherry::PH; unc-119(+)]* (Ziel *et al.* 2009).
LGIV: *qyIs10[lam-1::gfp; unc-119 (+)]* (Ziel *et al.* 2009).
LGV: *qyIs50[Pcdh-3>mCherry::moeABD, unc-119(+)]* (Ziel *et al.* 2009).
LGX: *dlg-1(mc103[dlg-1::gfp]*) (Vuong-Brender *et al.* 2017).

### Microfluidic Device Fabrication

Microfluidic devices were fabricated using standard soft- and photolithographic protocols (Xia *et al.* 1998). Briefly, master molds were fabricated on silicon wafers (Si-Wafer 4P0/>1/525±25/SSP/TTV<10, Siegert Wafer, Germany) using SU8 photoresist (GM1050 and GM1060, Gersteltec, Switzerland). Layers of different height were fabricated consecutively on the same wafer and aligned to each other using a mask aligner (UV-KUB3, Kloe, France). Masters of the fluidic layer for L1 devices were fabricated with heights of 3 and 8 μm, the L1-L4 device at heights of 5 and 12μm and L2-A devices at heights of 5 and 15 μm (for the feeding and worm sections, respectively). The master mold for the control layer was fabricated on a separate wafer at a single height of 20 μm. All wafers were treated with chlorotrimethyl silane (Sigma Aldrich, Switzerland) vapor prior to PDMS casting, to prevent adhesion of the PDMS to the wafer and SU8 structures. PDMS devices were assembled by first spin coating (750 rpm for 30 seconds) the fluidic wafer with a thin layer of PDMS (Elastosil RT601; ratio 20:1, Wacker, Germany). Simultaneously an approximately 4 mm thick layer of PDMS Elastosil RT601; ratio 20:1, Wacker, Germany) was cast on the control layer. Both PDMS layers were partially cured at 70°C. Once sufficiently cured the control layer was peeled from the wafer, cut to size and access holes punched (20G Catheter Punch, Syneo, USA). The control layer was then aligned to the fluidic layer using a stereomicroscope. Both layers were thermally bonded to one another overnight at 70°C (Unger *et al.* 2000). Bonded devices were peeled off the wafer, access holes punched (20G Catheter Punch, Syneo, USA) and bonded to cover glass (50×24 mm cover glass with selected thickness 0.17±0.01 mm, Hecht Assistant, Germany) using an air plasma (Zepto Plasma cleaner, Diener, Germany).

### Device Operation

First, the on-chip hydraulic valve was filled with DI water by applying a constant pressure to the dead-end valve channel (**Fig. S2b-c**). For this, a short piece of 1/16” tygon tubing (15157044, Fisher Scientific, Switzerland) filled with DI water was connected to both the microfluidic device and an off-chip solenoid valve, which in turn was connected to an external pressure source. The dead-end hydraulic channel was filled due to the air permeable nature of PDMS. Under pressure, air initially present in the channel was pushed into the PDMS material, allowing the channel to completely fill with liquid. Second, the device was filled with the bacteria suspension introduced using 1/32” Tygon tubing (15137044, Fisher Scientific, Switzerland) connected to a 1mL syringe (300-35-986, 30G blunt needle, Distrelec, Switzerland), and pressurized to remove all air bubbles from within the device (**Fig. S2d-f**). Finally, worms were introduced via the worm inlet. For worm-loading animals were loaded into a short piece of tubing (1/16” Tygon tubing, 15157044, Fisher Scientific, Switzerland) connected to a buffer-filled 1mL syringe (300-35-970, 23G blunt needle, Distrelec, Switzerland) (**Fig. S2h**). The tubing was connected to the worm inlet and worms were pushed into the device by gently applying and releasing pressure on the syringe plunger. Worms were pushed from the common inlet into individual channels of the trap array and halted at the end of the trap channels at the transition from the trap channel to the lower height food supply channel. Bacteria food was supplied at a flow rate of 1μL/hr, with an increase in flow rate to 100 μL/hr for 5 seconds every 30 minutes, to remove possibly accumulated debris (Aladdin AL-1000, WPI, USA). All worm loading was monitored at 10X magnification. Once channels were filled and suitable worms identified within the trap array, a high magnification, high NA objective was focused onto the device and image acquisition started. During image acquisition, the on-chip hydraulic valve was pressurized, effectively immobilizing worms within the trap channels. Pressure was released after image acquisition was completed leaving worms to move and feed freely.

### Image Acquisition

All images were acquired on an epifluorescence microscope (either a Ti-U, Nikon, Switzerland or DMRA2, Leica, Switzerland) equipped with an sCMOS camera (either a Prime 95B or Prime BSI, Photometrics, USA) and a fluorescence light source (either LedHUB, Omicron Laserage Laserprodukte GmbH, Germany or Spectra, Lumencor, USA) and brightfield LED (either pE-100wht, Coolled, UK or MCWHLP1, Thorlabs, USA). Z-stacks were acquired using a piezo objective drive (either Nano-F100, Mad City Labs, USA or MIPOS 100 SG, Piezosystems Jena, Germany). Images were acquired in up to three colors at a Z-spacing of 0.2-0.5 μm and a time interval of 15 minutes for up to 48 hours in all experiments. Smaller time intervals or longer overall image acquisition times may be chosen depending on the application. However, image acquisition may affect worm development if incompatible parameters are chosen. Images were acquired either using a 40X water immersion lens (HC PL APO 40x/1.10 W CORR CS, Leica, Switzerland), a 40X oil immersion lens (HCX PL APO 40X/1.30 OIL, Leica, Switzerland), a 60X water immersion lens (CFI Plan Apo VC 60XC WI, Nikon, Switzerland) or 60X oil immersion lens (CFI Plan Apo Lambda 60X Oil, Nikon, Switzerland). Image acquisition as well as actuation of the on-chip hydraulic valve was controlled through a custom written Matlab script (Matlab 2019b, Mathworks, USA) and a custom built microcontroller (Arduino Mega 2560, Arduino, Italy) for coordination of fluorescence and brightfield LEDs, piezo and camera.

### Image Deconvolution

Images were deconvolved either using the Huygens Deconvolution platform (SVI, Center for Microscopy and Image Analysis, University of Zürich) or the YacuDecu implementation of CUDA based Richardson Lucy deconvolution in Matlab.

### Bacterial Food Preparation

Worms were fed on-chip with *E. coli* NA22 grown overnight in LB medium at 37°C. 40 mL of saturated bacterial cultures were pelleted by centrifugation (3000 rcf for 2 minutes) and washed with fresh S-Basal buffer for a total of three washes. Finally, bacteria were pelleted and resuspended in 1mL fresh S-Basal, and 1 mL of bacterial suspension (50%) was mixed with 0.65 mL Optiprep (33%) (Sigma-Aldrich, Switzerland) for density matching of bacteria in suspension and 0.33 mL S-Basal supplemented with 1wt% Pluronic F127 (17%) (Sigma-Aldrich, Switzerland), a nonionic surfactant. Pluronic F127 was added to prevent bacteria aggregation in the low height section of the device and was found not to affect worm development at the chosen final concentration of 0.125%. Prior to filling the device, the bacteria suspension was passed through a 10 μm filter (pluriStrainer Mini 10 μm, PluriSelect, USA) to remove any large aggregates or debris. Bacteria food was prepared fresh for every experiment.

### Developmental Timing Assay

Developmental timing on chip was determined by manually identifying the first seam cell division in each larval stage as well as the completion of eversion in images acquired using a 40X magnification at 15-minute intervals for up to 48 hours. Worms were synchronized as described above, and for imaging in the L1 stage loaded directly onto the microfluidic device after overnight starvation. Animals imaged from late L1 to L4 stage or late L2 stage to adulthood were first grown on NGM plates for 16 hours (L1-L4 device) and 24 hours (L2-A device) after seeding. Developmental timing for free crawling worms was established on NGM plates seeded with *E. coli* NA22, the same bacterial strain used in on-chip experiments (to exclude any effect of food source on developmental timing). Worms were picked off the NGM plates every hour and the progression through seam cell division monitored. For each time-point on average 10-15 animals were imaged, resulting in the average developmental progression. For worms on plate developmental time was estimated as the median between the first time point at which animals of both stages are present on plate and the last time point at before all animals transitioned to the next stage.

### Seam Cell Lineage Assay

**S**ynchronized L1 populations were generated by bleaching gravid adult hermaphrodite animals and allowing the larvae to hatch overnight in S-Basal buffer. After overnight incubation, worms were filtered through a 10 μm filter and washed twice with fresh S-Basal. Worms were then loaded onto the L1 device and imaged at 15-minute intervals for a total of 48 hours using a 40X magnification objective. Seam cells were tracked manually using Fiji software (Schindelin *et al.* 2012) and the start of each cell division identified using the DLG-1::GFP marker.

### VPC induction Assay

Synchronized L1 larvae were initially grown on NGM plates and harvested approximately 16 hours after seeding (late L1 stage). Worms were washed and loaded as described, imaged using a 40X magnification, acquiring a full Z-stack every 30 minutes for a total of 48 hours. CFP-fluorescence intensity of the 1° fated VPCs and gonad length were measure manually over time using Fiji software (Schindelin *et al.* 2012). Measured maximum fluorescence intensity was normalized (using the maximum observed intensity in 2-cell or 4-cell stage) and background subtracted for comparison.

### Vulval Morphogenesis Assay

Worms were grown as described above with synchronized L1 larvae grown on NGM plates and harvested 24 hours after seeding. Worms were imaged at 60X magnification with the FOV set up, such that worms would remain in the available field-of-view throughout the experiment. For analysis, the developing vulva was cropped using a custom written Matlab script, followed by deconvolution of the acquired images. Measurements were taken manually at select time points using Fiji software (Schindelin *et al.* 2012).

### On-Chip RNAi

Prior to each experiment 20mL of LB medium supplemented with ampicillin (0.1mg/mL final concentration) were inoculated with bacteria from the stock LB plates and grown at 37°C over-night. Following this initial growth period additional 20mL of LB medium supplemented with IPTG (1mM final concentration) and ampicillin (0.1mg/mL) was added to the overnight culture and bacteria were grown for another 3 hours to induce dsRNA synthesis (Conte *et al.* 2015). After this second growth phase, bacteria were harvested and processed as described in the above section, resulting in the desired worm RNAi bacteria food. It should be noted that for RNAi a different bacterial strain is used (HT115). Therefore, the necessary concentration of Optiprep was adjusted from 0.65mL (33%) (stated above) to 0.59mL (30%), likewise the amount of S-Basal supplemented with Pluronic F127 was adjusted in compensation to 0.38mL and an additional 0.002mL of IPTG (1M stock concentration) was added.

Worms were initially grown on RNAi plates, produced by adding IPTG (1mM final concentration) and ampicillin (0.1mg/mL final concentration) to the standard NGM plate mixture and seeded with the respective RNAi bacterial strain. Bacteria for seeding were grown following the same scheme outlined above, starting from an overnight culture with ampicillin only, followed by a second 3-hour growth phase in the presence of IPTG. The bacterial culture was then concentrated to approximately 1/5^th^ of the initial volume and each NGM plate seeded with 0.3mL of concentrate. RNAi bacteria were allowed to grow on plate, at room temperature for at least 24 hours before seeding worms. Worms were prepared by bleaching of gravid adult animals as described above and grown on RNAi plates from L1 stage onwards, harvested approximately 24 hours after seeding and loaded into the microfluidic device as described. Images for both RNAi conditions were acquired at 15-minute intervals at 60X magnification for a total of 36 hours.

## Acknowledgements

We wish to thank the members of the Hajnal laboratory for critical discussion and comments on the manuscript. We are also grateful to the *C. elegans* Genetics Center CGC, which is funded by NIH Office of Research Infrastructure Programs (P40 OD010440) and J. Ahringer for RNAi clones. This work was supported by grants from the Swiss National Science Foundation no. 31003A-166580 to AH, the Swiss Cancer league no. 4377-02-2018 to AH and funding by ETH Zürich to AdM.

## Author contributions

SB conceived the method and designed experiments. SB performed all microfluidic experiments. SS performed NGM control experiments. SB and SS analyzed data. SB, AdM and AH wrote the manuscript.

